# Predator-prey feedback in a gyrfalcon-ptarmigan system?

**DOI:** 10.1101/220038

**Authors:** Frédéric Barraquand, Ólafur K. Nielsen

## Abstract

Specialist predators with oscillating dynamics are often strongly affected by the population dynamics of their prey, yet they do not always participate in a predator-prey cycle. Only those that exert strong population regulation of their prey do so. Inferring the strength and direction of the predator-prey coupling from time series therefore requires contrasting models with top-down versus bottom-up predator-prey dynamics. We examine such population-level coupling using multivariate autoregressive models. The models translate several hypotheses for the joint dynamics of population densities of the Icelandic gyrfalcon *Falco rusticolus*, and its prey, the rock ptarmigan *Lagopus muta*. The dynamics of both species are likely not only linked to each other but also to stochastic weather variables acting as confounding factors on the joint dynamics. The classical MAR(1) model, used most often in ecology, predicts that the times series exhibit predator-prey feedback (i.e., Granger causality): the predator helps to explain prey dynamics and the prey helps to explain predator dynamics. Weather, in the form of spring temperature, influences gyrfalcon population growth but not ptarmigan population growth, despite individual-level evidence that ptarmigan chicks can be strongly affected by weather. MAR(2) models, allowing for species to cycle independently from each other, further suggests alternative scenarios where a cyclic prey influence its predator but not the other way around; such bottom-up models produce a better fit but less realistic cross-correlation patterns. Simulations of MAR(1) and MAR(2) models further demonstrate that the top-down MAR(1) models are most likely to be misidentified as bottom-up dynamics than vice-versa. We therefore conclude that predator-prey feedback in the gyrfalcon-ptarmigan system is very likely, though bottom-up dynamics cannot be excluded with certainty. We finally discuss what sort of information is needed to advance the characterization of joint predator-prey dynamics in birds and other vertebrates.

## Introduction

Theoretical ecology predicts that among predators, specialists are the most likely to shape the dynamics of their prey (e.g. Andersson and Erlinge, 1977; Turchin and Hanski, 1997; Gilg et al., 2003). It has even been suggested that only specialist predators do exhibit multi-generation predator-prey population cycles (Murdoch et al., 2002), based on cycle periods in specialist versus generalist predators. Mechanistic modeling, however, disputes this particular point (Erbach et al., 2013). Testing more thoroughly this working theory with empirical data – the more specialized the predator, the higher the likelihood of a predator-prey cycle or more generally top-down prey regulation – would require to estimate the strength of predator-prey coupling in a number of real predator-prey systems, for which time series of both predator(s) and prey are available, preferably in the field. While the task may appear straightforward in theory, it is surprisingly difficult in practice. Cases of long-term monitoring including *both* specialized predators and their main prey, through extended periods of time, are indeed quite rare, especially in vertebrates. Two famous exceptions to the rule include the wolf-moose (*Canis lupus – Alces Alces*) system of Isle Royale (Vucetich et al., 2011), that has been followed for a century (although this study area is somewhat restricted for such wide-ranging species), and the celebrated cycle of the Canada snowshoe hare *Lepus americanus*, which interacts with the Canada lynx *Lynx canadensis* and other predators (Vik et al., 2008; Krebs et al., 2001). While there is a convincing array of evidence showing that lynx have a dynamical impact on hares (Vik et al., 2008), and wolf has an impact on moose (Vucetich et al., 2011), there is also evidence that weather and other drivers have often a strong forcing influence on prey dynamics (Vucetich and Peterson, 2004; Yan et al., 2013). Even in such strongly interacting systems that fascinate the imagination by demonstrating strong oscillations, it has been suggested that the presence of an ubiquitous external forcing hardly warrants to view such systems as a pair of autonomous coupled differential equations (Nisbet and Gurney, 1976; Barraquand et al., 2017), despite the pivotal role of autonomous and deterministic dynamical systems in ecological theory (McCann, 2011; Arditi and Ginzburg, 2012). Rather, real predator-prey systems are constantly buffeted by outside forces, be those climatic or biotic variables unaccounted for (i.e, other players in the interaction web). The study of Vik et al. (2008) reports at best around 55% of prey variance in log-densities explained by both prey and predator densities; this therefore leaves ample room for other factors to influence hare dynamics (Barraquand et al., 2017; see also Vucetich et al. 2011 on ungulate-wolf systems). In birds, contrasted feedback structures (bottom-up or top-down) were found in goshawk (*Accipiter gentilis*) – grouse dynamics, depending on the grouse species considered (Tornberg et al., 2013), with marked effects of weather forces. To gain a better appraisal of the strength of top-down regulation in the field, compared to other drivers of herbivore dynamics (see Sinclair, 2003, for a discussion in mammals), the list of predator-prey systems to which stochastic models of interacting populations are fitted to time series needs to increase. Having a number of reference predator-prey systems, whose dynamical structure (e.g., top-down vs. bottom-up) have been vetted by time series analysis, will also help evaluating how future food web models should be structured to obtain reliable quantitative predictions: what is the percentage of top-down links that should be allowed? Should bottom-up interaction coefficients generally be higher or lower than top-down? Should intra-specific density-dependence dominate? Should there be strong weather effects, strong or weak noise?

Our goal here is to contribute, using large-scale field data, to improving the understanding of predator-prey dynamics. We do this by fitting stochastic, statistically-driven predatorprey models to a presumably tightly coupled predator-prey pair, gyrfalcon *Falco rusticolus* and rock ptarmigan *Lagopus muta* in North-East (NE) Iceland. The gyrfalcon is a predator specialized on ptarmigan (rock ptarmigan and willow ptarmigan *Lagopus lagopus*) (Nielsen and Cade, 2017). In Iceland, the rock ptarmigan amounts to on average 72% by biomass of the gyrfalcon summer diet (range 52-86%) (Nielsen 1999). We combine detailed monitoring data from Iceland with multivariate autoregressive (MAR) modeling to infer the strength of trophic coupling in this system. Previous studies have computed autocorrelation functions to infer periodicity (Nielsen, 1999) and fitted autoregressive models on each species (Brynjarsdóttir et al., 2003), but this is to our knowledge the first time a MAR model is fitted to this dataset. MAR modeling, now an established technique within ecology (Ives et al., 2003; Hampton et al., 2013), has been largely developed in econometrics (Granger, 1969; Lütkepohl, 2005), where it is primarily used to establish causal relationships in the sense of prediction (i.e., a variable has causal influence if it helps improving predictions about the future, Granger 1969), which is the statistical philosophy that we adopt here.

## Material and methods

### Study area and design

The study area (5327 km^2^) in NE Iceland and survey methods used have been extensively detailed elsewhere (Nielsen, 1999, 2011) so we will remain brief. The gyrfalcon population is censused annually by visiting all known territories within the study area to determine predator occupancy (*n* = 83 territories). The number of territorial rock ptarmigan males is surveyed every spring (mostly in May) on 6 plots (total area 26.8 km^2^) within the general study area. The study started in 1981 and we use data for the period 1981-2014.

### Ecological variables

We consider two main variables, the occupancy rate of gyrfalcon territories, that was considered a good proxy for gyrfalcon population density, and mean density of territorial rock ptarmigan cocks on the 6 plots. Both variables are standardized in the statistical models.

We also consider weather variables that are known to potentially affect the dynamics of the two populations. We have selected 3 stations for the temperature (Akureyri, Mánárbakki, Grímsstaðir), and 6 stations for log-precipitation (Lerkihlíð, Mýri, Staðarhóll, Reykjahlíð, Mánárbakki, Grímsstaðir), all within or at the border of the study area and that have recordings from 1975 to now. The weather data was retrieved from the web site of the Icelandic Met Office (http://www.vedur.is/).

### Statistical models

Multivariate AutoRegressive (MAR) models have been used to assess the strength of predator-prey coupling (Ives et al., 2003; Vik et al., 2008). Let us denote the ln-transformed predator density *p_t_* = ln(*P_t_*) and *n_t_* = ln(*N_t_*) the ln-tranformed density of the prey; the log transformation is useful to transform log-normal into Gaussian noise. These ln-densities are then centered and stacked into a vector x_*t*_ = (*x*_1*t*_,*x*_2*t*_)′ = (*n_t_,p_t_*)′. The dynamics of the MAR(1) model, with one timelag, are then written as a forced recurrence equation (eq. 1),

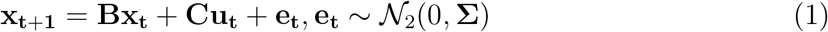

where B is an interaction matrix that characterizes the effects of net interactions on population growth of the 2 species, C describes the effect of environmental covariates **u**_*t*_ on the population growth rates of predator and prey, and e_t_ is a Gaussian bivariate noise term.

We considered both a model without interactions where 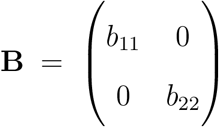, hereafter referred to as the null MAR(1) model, and a model with full interaction matrix, B = 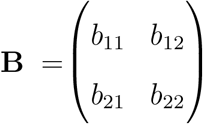. Both models were considered without (C = 0) and with environmental forcing (C ≠ 0). The weather variables that we stacked within **u**_*t*_ are delayed: the predator population is believed to be affected by weather 5 years before, because recruits enter the adult population at the age of 4 years (average time to maturity), while the prey population is affected by the weather of the year (between *t* and *t* + 1) or that of the preceding year (between *t* – 1 and *t*). We considered models with temperature effects, log(precipitation) effects, or both.

Several model fitting techniques have been considered in preliminary explorations (MCMC using JAGS within `R`, least squares for vector autoregressive models in R package **`vars`**, simple independent linear autoregressive models using `lm()` in R). Maximum likelihood estimation using the MARSS package (Holmes et al., 2012) and the EM algorithm was finally chosen because it allowed to easily perform model selection for contrasted interactions matrices (i.e., setting some interactions to zero). All algorithms gave however similar model estimates (see Appendix S1).

We then considered more complex MAR(2) models that are able to allow for both populations to cycle independently, because each univariate AR(2) component can model long cycles (≈ 7 to 10 years cycles, like those observed in the field). Selection of the optimal lag *p* in MAR(p) model using a variety of model information theoretic criteria (see code in **https://github.com/fbarraquand/GyrfalconPtarmigan_MAR**) in suggested an optimal lag order of 2 (BIC, HQ) or 3 (AIC, FPE). Because 2 lags are enough to model independently cycling populations of period up to 10 years and more (Royama, 1992), and MAR(2) models are already parameter-rich, we considered a maximum of 2 lags in MAR models. The MAR(2) model can be written as

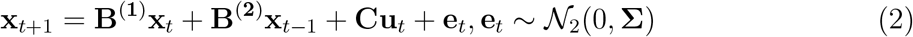

The independent cycling model has diagonal matrices B^(1)^ and B^(2)^. The full model has interaction matrices 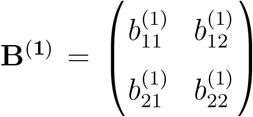 and 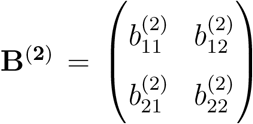. To model also an asymmetric and nonreciprocal effect from the cyclic prey to its predator, we used the following interaction matrices 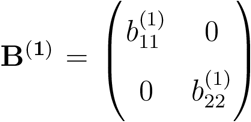 and 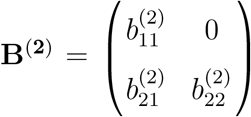. The model was named ‘bottom-up’, in order to designate a predator dynamics driven by that of its cyclic prey.

## Results

### MAR(1) model results

#### Models without environmental covariates

The predator-prey time series and the MAR(1) model one-step ahead predictions are presented in Fig. 1, while Table 1 shows the MAR(1) model fitted parameters. All **B** coefficients are found to be significantly different from zero, with commensurate strengths of predator → prey (*b*_12_) and prey → predator (*b*_21_) interaction. There is therefore consistently negative effect of predator on prey and a consistently positive effect of prey on predator. Note that a clear-cut sign was not obligatory, given those are net interaction coefficients, blending several ecological processes, e.g., direct and indirect predation effects, into one number.

**Figure 1:**
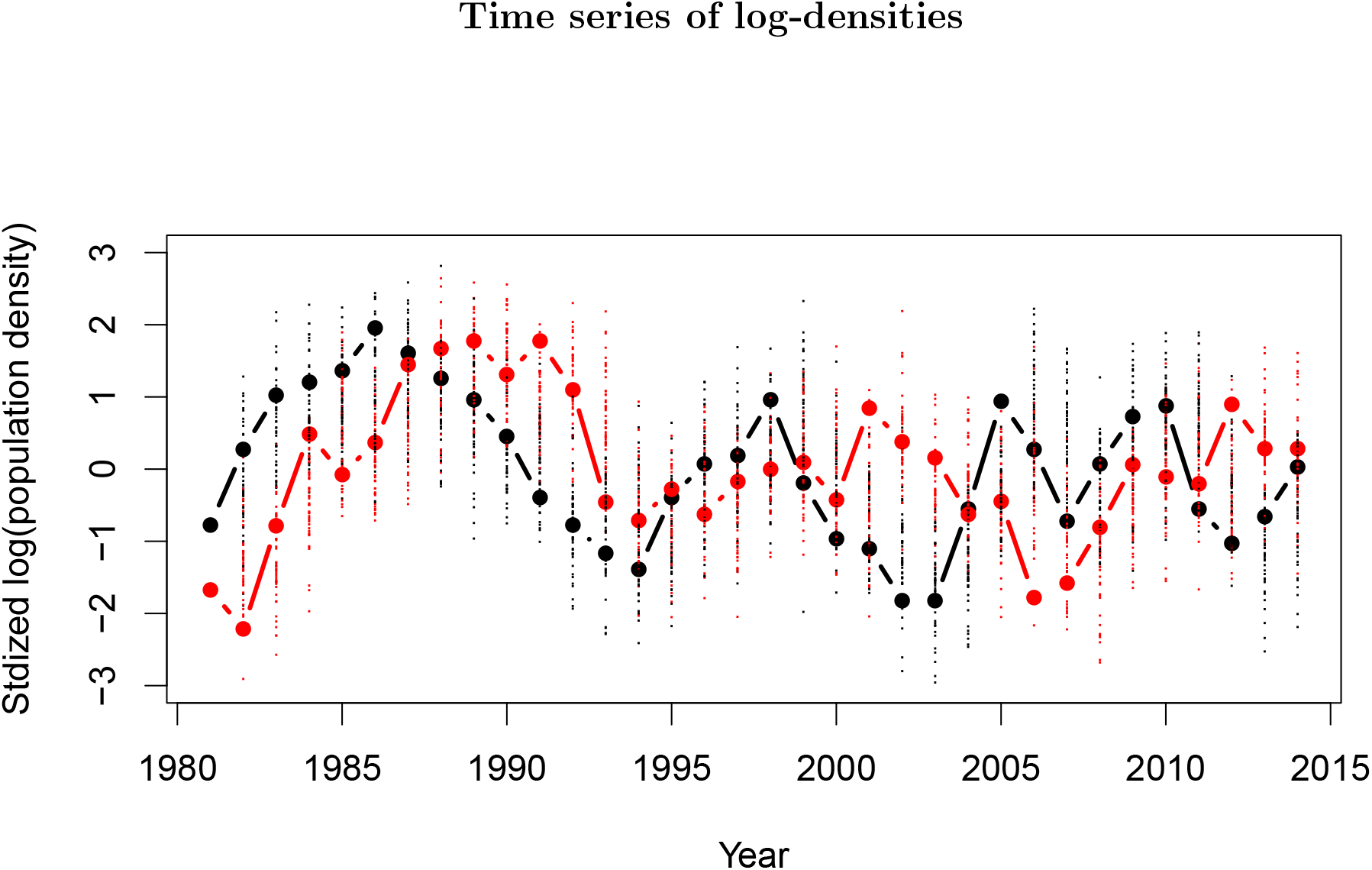
Time series of gyrfalcon (red) and rock ptarmigan (black) standardized log-densities in NE Iceland, and their corresponding one-step ahead predictions under the best-fitted, full interaction matrix MAR(1) model. 100 model simulations one step ahead are plotted, for each year, as small points – red for predator and black for prey.

**Table 1:**
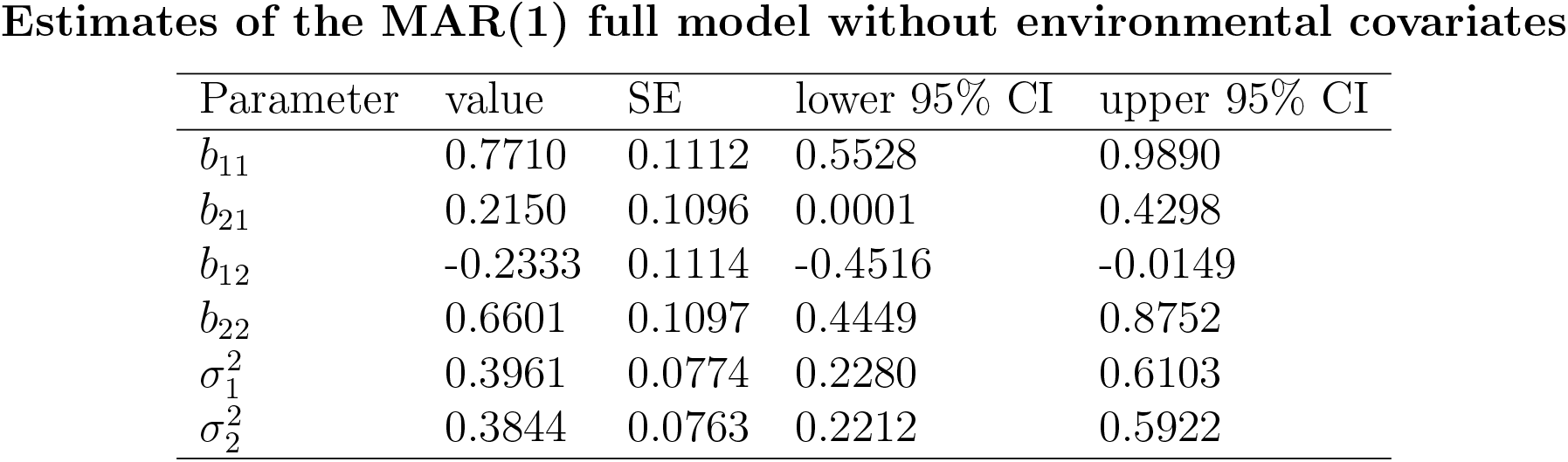
The off-diagonal interaction coefficients are statistically significant at a 95% level. Similar results are obtained for non-diagonal Σ (not shown); for parsimony we use a diagonal error matrix.

**Table 2:** MAR(1) ‘null’ indicates a diagonal B matrix while MAR(1) ‘full’ indicates a full 2x2 interaction matrix. Models including temperature effects on growth rates (third row and below) effects on growth rates take the form x_t+1_ = Bx_t_ + Cu_t_ + e_t_, e_t_ ∼ N_2_(0, Σ). Here the environmental vector is u_t_ = (*T*_*t-lp*+1_, *R*_*t-lP*+1_,*T*_*t-l_G_*+1_, *R*_*t-l_G_*+1_)^*T*^, with *T* the temperature and *R* log-precipitation. There is a time-lag *l_P_* for the ptarmigan (0 or 1 year) and *l_G_* = 5 (always) for the gyrfalcon: weather is clearly expected to have such delayed effects on the gyrfalcon counts because of age structure. April weather is considered for gyrfalcon as it is the critical period for reproduction, and it is always included in models from row 3 and below. Models from rows 3 to 6 considered May temperature for ptarmigan, log(precipitation), or both. The models of rows 7 and 8 considered instead July and June temperatures as environmental variables for ptarmigan.

| Model type | LogLik. | AIC | AICc | BIC |
| --- | --- | --- | --- | --- |
| MAR(1) null | -70.01 | 148.0 | 148.7 | 154.1 |
| MAR(1) full | -66.14 | 144.3 | 145.7 | 153.4 |
| MAR(1) full + May temperature year $t + 1$ | -63.98 | 144.0 | 146.4 | 156.2 |
| MAR(1) null + May temperature year $t + 1$ | -67.03 | 146.1 | 147.4 | 155.2 |
| MAR(1) full + May temp. of year $t$ | -63.94 | 143.9 | 146.3 | 156.1 |
| MAR(1) full + May log(precipitation) of $t$ | -64.89 | 145.8 | 148.2 | 158.0 |
| MAR(1) full + May temp + log(precipitation) | -62.81 | 145.6 | 149.5 | 160.9 |
| MAR(1) full + July temperature year $t$ | -61.95 | 143.9 | 147.8 | 159.2 |
| MAR(1) full + June temperature year $t$ | -63.79 | 147.6 | 151.4 | 162.8 |

Based on the comparison of AICc and BIC between the full 2 x 2 interaction matrix and the diagonal matrix model (null model), the full model was favored (Table 2). The model with environmental covariates did not lead to substantially better fit or very different biotic interaction parameters (Table 2) than the full model, and the weather effects were not consistent (Table 3), save for those of delayed April temperature on predator growth (see below).

**Table 3:** Species 1 is ptarmigan and species 2 gyrfalcon. May variables only affect species 1 while April variables, delayed by 5 years (we model the effect of variables at *t* – 4 on growth between *t* and *t* + 1), affect only species 2’s population growth.

| Parameter | value | SE | lower 95% CI | upper 95% CI |
| --- | --- | --- | --- | --- |
| $b_{11}$ | 0.7475 | 0.1071 | 0.5376 | 0.9575 |
| $b_{21}$ | 0.2031 | 0.1044 | -0.0014 | 0.4077 |
| $b_{12}$ | -0.1984 | 0.1135 | -0.4210 | 0.0241 |
| $b_{22}$ | 0.7021 | 0.1043 | 0.4977 | 0.9065 |
| temperature May$_{t+1}$ | -0.0853 | 0.1142 | -0.3092 | 0.1385 |
| precipitation May$_{t+1}$ | -0.1596 | 0.1091 | -0.3735 | 0.0541 |
| temperature April$_{t-4}$ | 0.2072 | 0.1061 | -0.0008 | 0.4153 |
| precipitation April$_{t-4}$ | -0.0558 | 0.1068 | -0.2652 | 0.1535 |
| $\sigma_1^2$ | 0.3653 | 0.0743 | 0.2104 | 0.5628 |
| $\sigma_2^2$ | 0.3418 | 0.0733 | 0.1943 | 0.5307 |

Additional Granger causality testing using the MAR(1) model revealed a two-way reciprocal feedback, though the Wald test was weakly statistically significant (at the 0.1 level) due to the low sample size (i.e., not by ecological standards but compared to other fields using time series analysis such as econometrics). While accounting for the relative shortness of ecological time series, the MAR(1) model therefore strongly suggests a reciprocal predator-prey coupling of the ptarmigan and gyrfalcon populations.

#### Models with environmental covariates

The addition of environmental covariates did not improve significantly model fit (Table 2). The coefficients were mostly non-significant, as illustrated by the model including both temperature and log(precipitation) (0 is included within CIs for environmental **C** matrix coefficients, Table 3). The model with both precipitation and temperature was deemed over-parameterized by the information criteria. It is likely that an effect of 5-year delayed temperature on predator growth is present as this effect was found positive, large and nearly statistically significant at 95% (Table 3). Although this weather effect on the predator does not seem to improve significantly the predictive ability of the model. The effect of temperature in May_*t*+1_ (May of the year) on ptarmigan growth, by contrast, is both not statistically different from zero and of unexpected sign (negative here, while positive temperature usually have positive effects on the ptarmigan chicks, Nielsen et al. 2004). The effect of rain in May_*t*+1_ on ptarmigan population growth was negative and relatively strong, but not statistically significant at 95% (point estimate −0.1596, 95%CI: [−0.3735; 0.0541]). It is therefore possible that such a negative effect is present, but it does not appear clearly with the current dataset.

We also fitted models where winter weather affects ptarmigan growth (Appendix S2), to test the idea that the survival of first-year chick might be lower in harsher winters, but again we did not find consistent effects of weather on ptarmigan growth.

### MAR(2) model results

The MAR(2) models showed uniformly better fit than the MAR(1) models (Table 4). Note that, in order to make this comparison, we re-fitted the MAR(1) model with one less year to compare MAR(1) and MAR(2) models with an equal number of points, as any difference in data can strongly affect AIC and BIC values. The MAR(2) model with independent populations (i.e., diagonal interaction matrices **B**^(1)^ and **B**^(2)^) and the bottom-up predator-prey model (see Methods), assuming an independently cycling prey and a predator whose dynamics is forced by its prey, were the better-ranking models (Table 4).

**Table 4:** Comparison of model selection criteria for MAR(1) and MAR(2) models with different structures. See Methods for definitions. The MAR(2) null + temperature uses April temperature with a 5 year delay, which affects the predator only – this model adds temperature to the list of potential drivers for predator dynamics, as it was found marginally significant in previous MAR(1) analyses.

| Model type | logLik. | AIC | AICc | BIC |
| --- | --- | --- | --- | --- |
| MAR(1) null | -67.57 | 143.1 | 143.8 | 149.1 |
| MAR(1) full | -64.41 | 140.8 | 142.3 | 149.8 |
| MAR(2) full | -54.78 | 129.6 | 133.6 | 144.5 |
| MAR(2) full | -57.91 | 129.8 | 131.8 | 140.3 |
| MAR(2) null | -58.78 | 129.6 | 131.0 | 138.5 |
| MAR(2) null + temperature | -56.95 | 129.9 | 132.4 | 141.9 |

Because AIC and BIC assess only one aspect of statistical model quality, the trade-off between model parsimony and fit, we also present the results of simulations of the models (Fig. 2). The examination of time series plots is however difficult because the simulated time series are relatively short and noisy.

**Figure 2:**
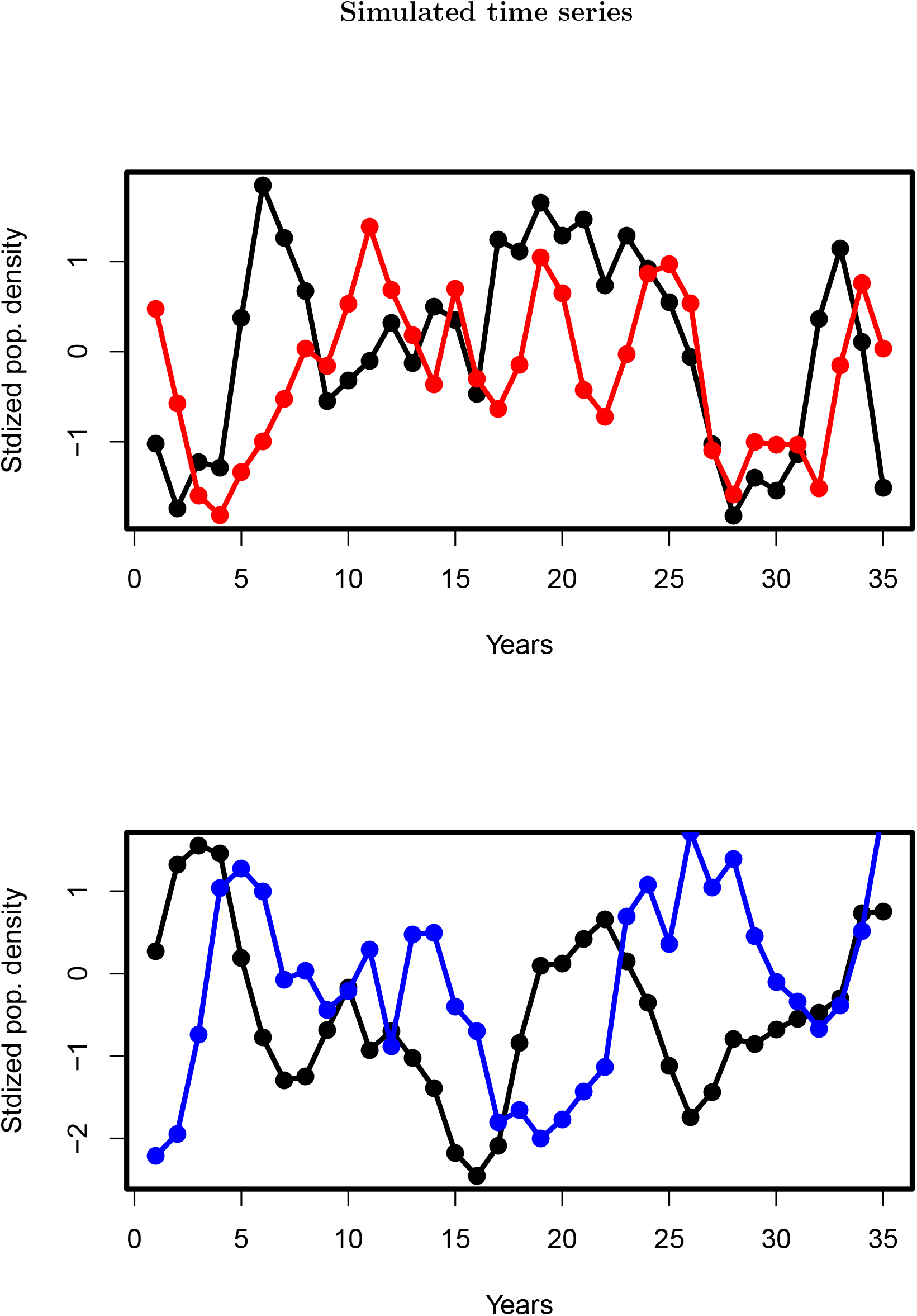
Time series of predator (gyrfalcon) and prey (rock ptarmigan) log-densities, simulated for 35 years from the same starting conditions as the data, for the full MAR(1) model (top panel, predator in red) and the MAR(2) ‘bottom-up’ model (bottom panel, predator in blue).

We therefore simulated 100 datasets using the fitted models (Fig. 3) and examined their cross-correlations, these show that MAR(1) and MAR(2) models with reciprocal predator-prey feedback (full interaction matrices) outperform both the bottom-up model (medium reproduction of the the cross-correlation pattern) and the null model (no reproduction of the cross-correlation pattern).

**Figure 3:**
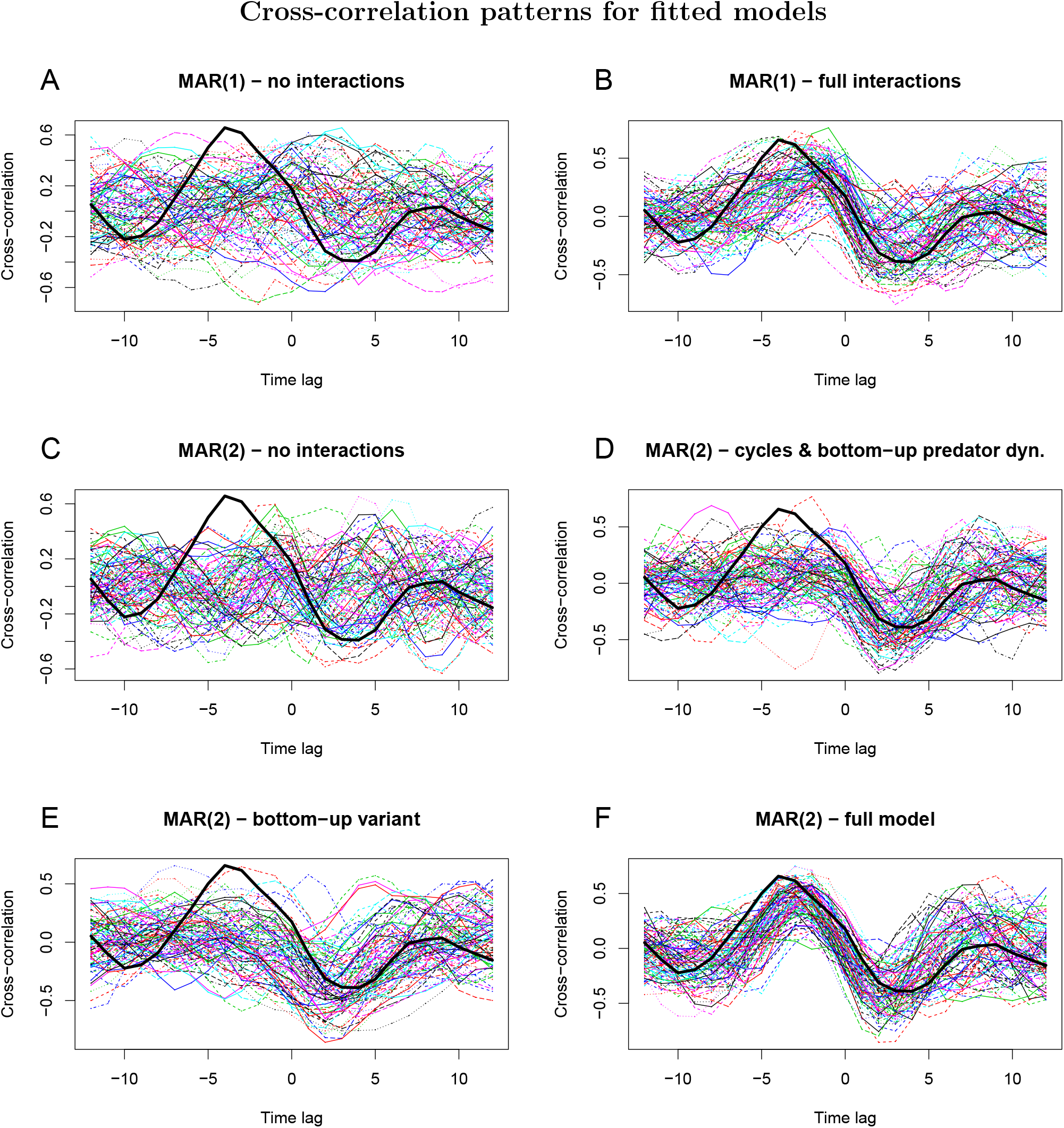
Cross-correlation functions (CCFs) for the fitted models (A to F). Each thin line corresponds to one simulation of the fitted model, within each panel. A and B show MAR(1) models, without and with interactions; while C to F show the CCFs of simulated MAR(2) models, without interactions (C), with only bottom-up interactions (D), bottom-up without predator regulation with a delay (E), and (D) full MAR(2) model. The cross-correlation for the real data is highlighted as a thick black line in all panels.

## Discussion

The percentage of explained variance in log-abundances by the MAR(1) predator-prey model was about 60%; hence similar to the lynx-hare example of Vik et al. (2008). The full 2x2 interaction matrix in a MAR(1) framework provided a better model than a diagonal matrix, meaning there was causality (or *feedback*) between prey and predator dynamics in the sense of Granger (1969): the addition of the predator and prey variables reduced the residual variances of the time series models for the prey and predator, respectively.

Weather (i.e., April temperature 5 years lagged) was found to influence predator dynamics, revealing an influence of weather on gyrfalcon reproduction, which takes several years to affect the growth of the adult segment of the population. However, prey population growth was not affected by any of the weather covariates considered, neither in spring nor in winter. This is a surprising find, which we discuss below.

If we had stopped the analyses to the MAR(1) model, which is customary in ecology (e.g. Ives et al., 2003; Hampton et al., 2013), we would have concluded unequivocally to a strong coupling between predator and prey (Table 2). However, another reasonable hypothesis was that both species – the prey especially – could cycle independently (see e.g., Dobson and Hudson, 1992, for a host-parasite modeling study in a similar prey species). Contrasting such hypotheses required to formulate a MAR(*p*) model with *p* = 2 timelags (according to BIC, the optimal lag order was 2; 3 according to AIC). MAR(2) models were therefore found to realize a better trade-off between parsimony and fit than MAR(1) models (Table 4, for more information criteria see additional analyses^1^). While the model with independently cycling populations fitted the data well, it produced unrealistic dynamics (i.e., no cross-correlation, Fig. 3). The bottom-up predator-prey model, where the prey influences the predator but not the other way around, provided both a good fit and relatively realistic dynamics, though not as much as the models including reciprocal feedback (full-matrix MAR(1) and MAR(2) models). The bottom-up scenario could correspond, for example, to a case where the predator dynamics are driven by its prey, but prey dynamics are themselves driven by an interaction with a parasite (see Stenkewitz et al., 2016, for an appreciation of host-parasite dynamics in Iceland rock ptarmigan). The bottom-up scenario fitted the data better in terms of trade-off between parsimony and fit, but predicted the cross-correlation pattern worse. Hence with the currently available information, both scenarios must be considered plausible. Further simulation results do help, however, to interpret a little better which scenario is the most likely (see below).

This absence of conclusion on mechanisms, given the length of the survey, may appear at first sight distressing. However, from the perspective of time series analyses, even 35 years is very short. In fact, in his authoritative book on multivariate time series modeling, Lütkepohl (2005) shows that it can be hard to recover the simulated lag order of such simple 2 x 2 MAR(1) and MAR(2) models. Specifically, Lütkepohl (2005) simulated a bivariate MAR(2) model with a time series length *n* = 30, fitted MAR(p) models up to order *p* =6, and found only 32% of correctly classified simulations as *p* = 2 using AIC, with 42% classified as *p* =1 (p. 155 in Lütkepohl 2005). Using BIC, he found even 80% misclassified as MAR(1). Different model selection criteria give different answers, and the BIC tends to be most conservative, but the baseline is that for *T* = 30, selection according to information criteria only gives inconsistent answers, while in most cases (70% for AIC) the right model order was found for *n* =100. Results were overall better with simulated bivariate MAR(1) models (p. 156 in Lütkepohl 2005), where all model selection criteria were able to pinpoint the correct lag order at 90%. The results, however, are likely to be model-structure and model-parameter specific; therefore we performed such analysis for the models that we fitted. Specifically, we performed 1000 simulations of the fitted MAR(1) full (F) model and MAR(2) bottom-up (BU) model. We fitted the MAR(1) F model to both MAR(1) F and MAR(2) BU simulations, and then fitted the MAR(2) BU to both MAR(1) F and MAR(2) BU simulations. This allowed to compute the percentage of correctly ascribed scenarios, on the basis of information criteria (AIC, AICc, BIC) for each simulated model (Appendix S3). We found that for *n* = 35, a simulated MAR(1) F was recovered respectively 52%, 56%, 64% for AIC, AICc, and BIC, respectively. By contrast, a simulated MAR(2) BU model was recovered 98%, 98% and 97%. With the current length of our dataset, the important message from these simulations is that based on AIC or BIC, we are much more likely to mistake a fully interacting predator-prey system for a bottom-up system than the reverse. These percentages were all very close to 90% and 100% in the case of *n* = 100 (Appendix S3).

From our simulation experiments, we can derive three lessons. First, from an ecological viewpoint, given that the full interaction MAR(1) model both predicts the cross-correlation pattern better and is the most likely to be misidentified as MAR(2) BU, we should not give too much weight to the better (lower) AIC and BIC scores of the MAR(2) BU model. It is fairly likely that top-down prey regulation and therefore reciprocal predator-prey feedback is at work here. Second, from a more statistical viewpoint, it is informative to notice that whether MAR(1) or MAR(2) models are better identified is parameter-specific: sometimes a MAR(1) model will be more likely to be correctly classified (as in the simulation study of Lütkepohl, 2005), sometimes a MAR(2) will (our case study). Therefore, whether an ecological scenario is better identified than another one from time series is likely to be context-specific, and not simply dependent on the lag order. The corollary being that new simulations from MAR(p) or other time series models will be required for each new ecological case study, in order to see which scenarios are the most likely to be misidentified. Third, we found, in agreement with Lütkepohl (2005), that time series of around 100 points are needed to allow for fairly reliable inference of top-down vs bottom-up dynamics in systems of 2 cyclic species (less points might be required for species with simpler dynamics).

Given that the data presented here is collected once a year for the most part, and that it is not feasible to census the population much more frequently with current means (other technologies would be necessary, such as camera traps or DNA-based evidence), it is unlikely that we will get the time series near 100 years within acceptable time frames for management of both populations (i.e., conservation of the gyrfalcon and sustainable hunting management of ptarmigan). Therefore, differentiating unequivocally between the bottom-up and predator-prey feedback scenarios will likely require other type of models and data. We still view the MAR(p) approach as useful, however, as a means to delineate likely scenarios to investigate further, and check for important abiotic drivers that need to be considered. Why the ptarmigan population growth is not affected – at the NE Iceland scale – by weather variables is also a puzzling question for further study.

Mechanistic modeling might help further understand the effect of drivers on ptarmigan dynamics during certain phases of the cycle. For instance, are ptarmigan declines mainly driven by its predator (i.e., the gyrfalcon is responsible in large part for the declines) or mainly by other causes such as parasites? Rough estimates of predation demonstrate why this question is intrinsically difficult. Around 100 adult predator pairs can be found near peak abundance on the NE Iceland ptarmigan management zone, and to these correspond about 100 000 ptarmigan individuals at best (Sturludóttir, 2015). One might think, given these numbers, that the predators are unlikely to make their prey decline. Ptarmigan, however, have a slow, long-period cycle (Fig. 1). Therefore, they decrease at worst by ≈ 20 000 in a single year. A quick division indicates that about 200 would have to be eaten during the year by a predator pair for such a decrease to occur – assuming, as a first approximation, that increases in the ptarmigan population due to reproduction are offset by other causes of death than predation. This quantity, 200 kills a year, is an order of magnitude that represents a fairly high yet doable consumption by a predator pair. This might be tested further by fitting more mechanistic predator-prey models.

Although we currently do not possess all the information necessary to parameterize mechanistic predator-prey or host-parasite models, we suggest a few directions. First, we have a rather imperfect knowledge of the predator population, especially its non-territorial segment (Nielsen, 2011). Non-territorial floaters can indeed be rather numerous in both real raptor populations (Katzner et al., 2011) and parameterized bird population models (Barraquand et al., 2014). Floater numbers could therefore change our perception of predator impacts on prey dynamics (i.e., the predator population might increase by half or more). Demographic modeling of the predator population and its various life stages is therefore in order – we are currently examining CMR data and hoping for DNA-based information. Second, the current models have shown that the host-parasite hypothesis for the ptarmigan dynamics (see Stenkewitz et al., 2016) needs to be examined. We therefore need to know more about the parasite loads and their potential impact on ptarmigan population growth. Third, there are spatial aspects in the dynamics of gyrfalcon and ptarmigan that we have not tackled. It is plausible, for instance, that the weather does not affect ptarmigan growth at the scale of NE Iceland, using averaged state variables and covariables, and yet that weather locally affects the survival of ptarmigan chicks, as additional data seem to suggest: Nielsen et al. (2004) found that mean windspeed and mean precipitation in June-July explained a considerable part of the variance in chick production.

## Conclusion

Our results have implications for other studies on birds and more generally vertebrates with relatively slow life histories (compared to e.g., plankton sampled many times a year). Using long time series by ecological standards (34 years), we found some evidence of reciprocal predator-prey feedback in this cyclic predator-prey system, without being able to exclude nonetheless more bottom-up predator-prey dynamics. MAR(*p*) models with *p* =1,2 described well this system as a forced oscillator, although the unexplained noise was generally stronger than weather effects, which may point to other important biotic factors driving the dynamics, such as parasites. Simulations of the fitted models revealed than unequivocal inference of bottom-up versus reciprocal predator-prey coupling (i.e., including top-down predator influence on prey) would require about a century of time series data. We therefore think that additional demographic data (e.g., through capture-recapture, genetics,…) should always be considered in conjunction to counts taken once or twice a year, if one of the goals of a monitoring study is to infer interactions between the populations of different species.

## Acknowledgments

Financial support for 1981–1993 was received from the National Geographic Society, the Icelandic Science Fund, The Peregrine Fund, Inc., the Andrew Mellon Foundation, the E. Alexander Bergstrom Memorial Research Fund, and the Arctic Institute of North America. For 1986 and 1994–2014, this research was funded by the Icelandic Institute of Natural History. FB was supported by Labex COTE (ANR-10-LABX-45). The Lake Mývatn Research Station provided field facilities. Field assistance was provided by J.Ó. Hilmarsson, G. Þráinsson, I. Petersen, H. Bárðarson, Ó. Einarsson, E.Ó. Þorleifsson, Ó.H. Nielsen, A.Ö. SnæÞórsson, Þ.Þ. Björnsson, S. Nielsen. Many more people have helped in various ways with the field studies. These studies started out as OKN Ph.D. at Cornell University under the supervision of T.J. Cade.

## Footnotes

1 https://github.com/fbarraquand/GyrfalconPtarmigan_MAR The repository will be made publicly available upon acceptance.

## Supplementary Information

### Appendix S1 – Alternative model fitting

**Table S1:**
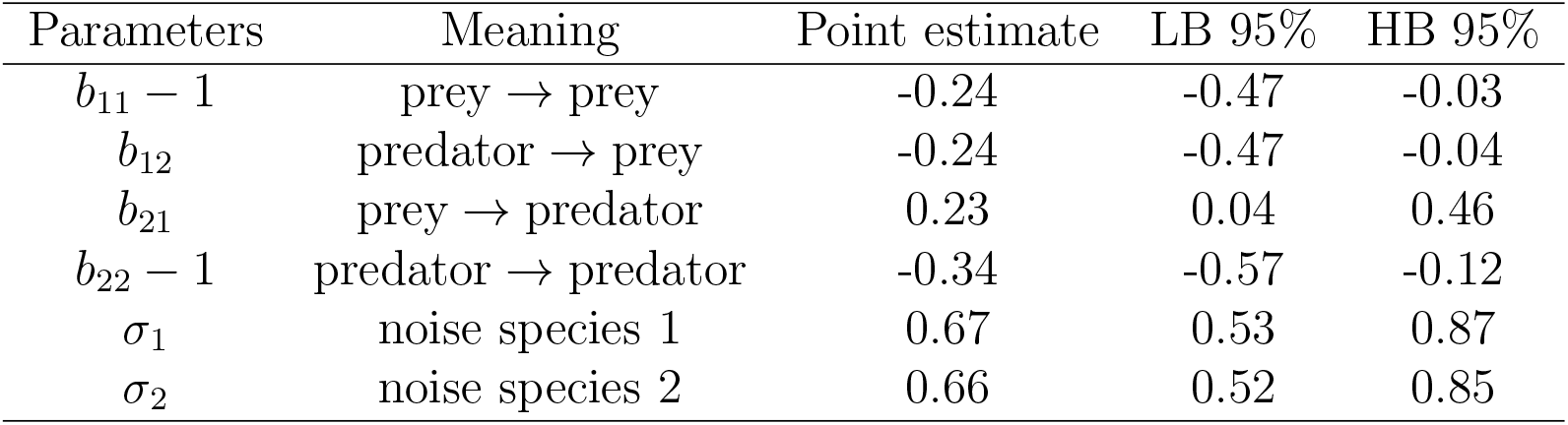
Estimates of the full MAR(1) model using JAGS, for comparison with the results of the main text.

### Appendix S2 – Effects of winter weather

We tested the effect of winter weather by introducing new winter weather variables into the MAR(1) models:

- The mean winter temperature from December to March
- The average of log(precipitation) over the same period

We also considered minimum temperature but this did not alter the following results.

The two above mentioned winter weather variables were inserted in place of spring weather variables for ptarmigan into a MAR(1) model. The estimated parameters are reproduced in Table S1 and the Information Criteria, with previous models for comparison, in Table S2. None of the models are able to significantly improve the fit, although it is possible that a weakly statistically significant effect of winter temperature exists.

**Table S1:**
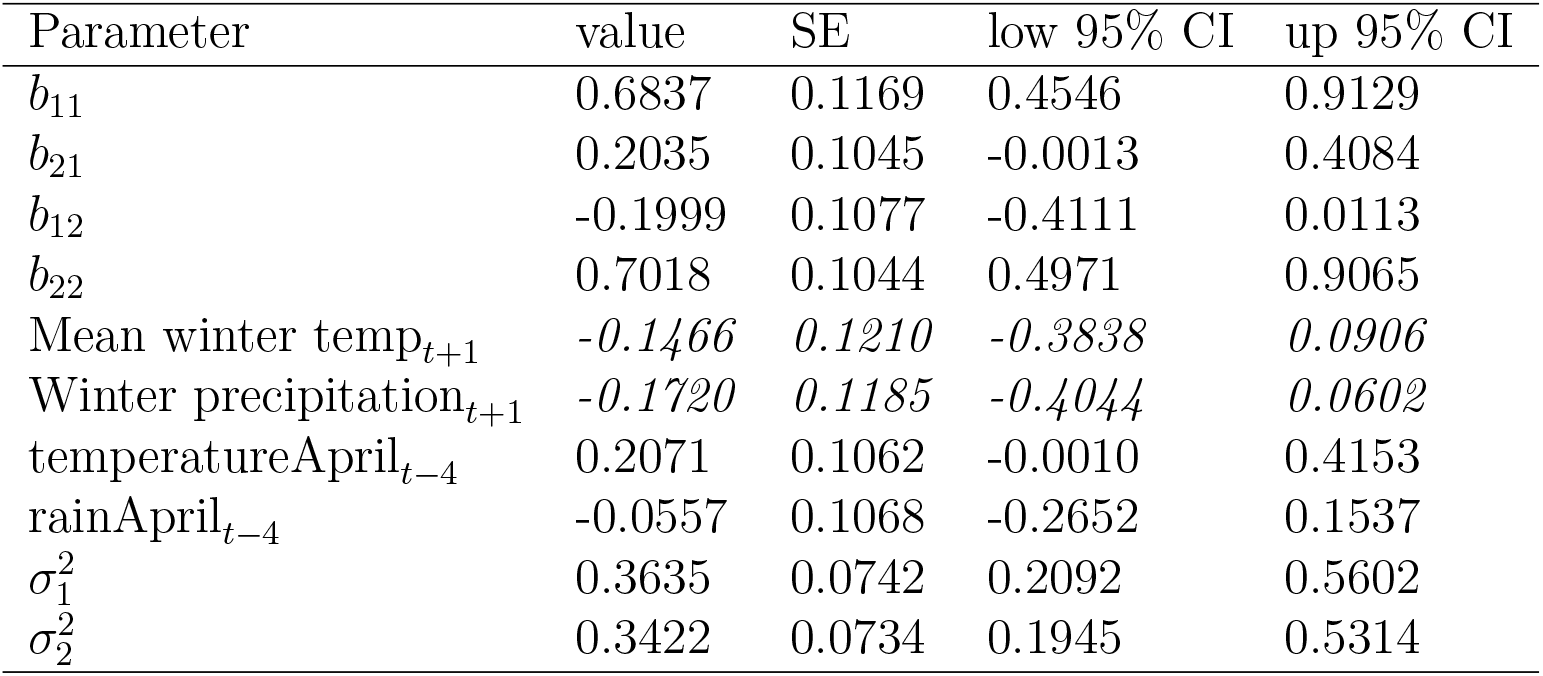
Coefficients for biotic and abiotic effects on population growth. Species 1 is ptarmigan and species 2 gyrfalcon. Winter variables only affect species 1 while April variables, delayed by 5 years (we model the effect of variables at *t* – 4 on growth between t and t + 1), affect only species 2’s population growth. *Effects of winter variables are depicted in italics*.

| Parameter | value | SE | low 95% CI | up 95% CI |
| --- | --- | --- | --- | --- |
| $b_{11}$ | 0.6837 | 0.1169 | 0.4546 | 0.9129 |
| $b_{21}$ | 0.2035 | 0.1045 | -0.0013 | 0.4084 |
| $b_{12}$ | -0.1999 | 0.1077 | -0.4111 | 0.0113 |
| $b_{22}$ | 0.7018 | 0.1044 | 0.4971 | 0.9065 |
| Mean winter temp$_{t+1}$ | *-0.1466* | *0.1210* | *-0.3838* | *0.0906* |
| Winter precipitation$_{t+1}$ | *-0.1720* | *0.1185* | *-0.4044* | *0.0602* |
| temperatureApril$_{t-4}$ | 0.2071 | 0.1062 | -0.0010 | 0.4153 |
| rainApril$_{t-4}$ | -0.0557 | 0.1068 | -0.2652 | 0.1537 |
| $\sigma_1^2$ | 0.3635 | 0.0742 | 0.2092 | 0.5602 |
| $\sigma_2^2$ | 0.3422 | 0.0734 | 0.1945 | 0.5314 |

**Table S2:**
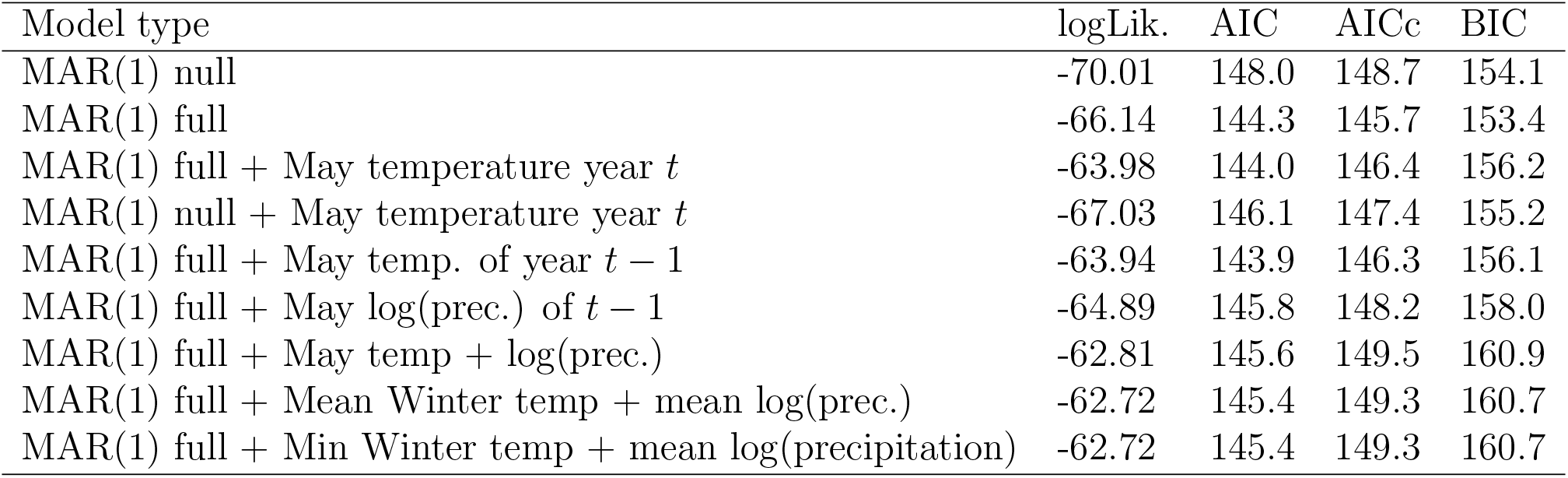
Comparison of model selection criteria for MAR(1) models. MAR(1) ‘null’ indicates a diagonal **B** matrix while MAR(1) ‘full’ indicates a full 2x2 interaction matrix. Models including temperature (third row and below) effects on growth rates take the form x_t+1_ = a + Bx_t_ + Cu_t_ + e_t_, e_t_ ∼ N_2_(0, Σ). Here the environmental vector u_t_ = (*T_t-lP_, R_t-lP_, T_t-lG_, R_t-lG_*)^T^, with *T* the temperature and *R* log-rainfall. There is a timelag *lp* for the ptarmigan (0 or 1 year, depending on the month) and *l_G_* = 5 for the gyrfalcon. IC scores for the two winter models are depicted on the last two rows. prec. = precipitation (rain and snow).

| Model type | logLik. | AIC | AICc | BIC |
| --- | --- | --- | --- | --- |
| MAR(1) null | -70.01 | 148.0 | 148.7 | 154.1 |
| MAR(1) full | -66.14 | 144.3 | 145.7 | 153.4 |
| MAR(1) full + May temperature year $t$ | -63.98 | 144.0 | 146.4 | 156.2 |
| MAR(1) null + May temperature year $t$ | -67.03 | 146.1 | 147.4 | 155.2 |
| MAR(1) full + May temp. of year $t - 1$ | -63.94 | 143.9 | 146.3 | 156.1 |
| MAR(1) full + May log(prec.) of $t - 1$ | -64.89 | 145.8 | 148.2 | 158.0 |
| MAR(1) full + May temp + log(prec.) | -62.81 | 145.6 | 149.5 | 160.9 |
| MAR(1) full + Mean Winter temp + mean log(prec.) | -62.72 | 145.4 | 149.3 | 160.7 |
| MAR(1) full + Min Winter temp + mean log(precipitation) | -62.72 | 145.4 | 149.3 | 160.7 |

### Appendix S3 – Simulation results

**Table S1:** Frequency of correct identification of MAR(1) full and MAR(2) bottom-up models for two time series lengths.

| Simulated model | AIC | AICc | BIC |
| --- | --- | --- | --- |
| $n = 35$ | 0.52 | 0.56 | 0.64 |
| $n = 100$ | 0.98 | 0.98 | 0.97 |

